# Ratiometric control of two microbial populations via a dual chamber bioreactor

**DOI:** 10.1101/2024.03.08.584056

**Authors:** Sara Maria Brancato, Davide Salzano, Davide Fiore, Giovanni Russo, Mario di Bernardo

## Abstract

Maintaining stable coexistence in microbial consortia, particularly when one species grows faster than another (i.e. the species are non-complementary), poses significant challenges. We introduce a novel control architecture that employs two bioreactors. In this system, the slower-growing species is cultivated separately before being introduced into the main mixing chamber. We analyze the open-loop dynamics of this setup and propose a switching feedback mechanism that controls the dilution rates to ensure robust regulation of population density and composition within the microbial consortium. Validated *in silico* using parameters from real experiments, our approach demonstrates effective and robust maintenance of microbial balance across various strains without requiring genetic modifications.

## I. INTRODUCTION

Microbes in nature are often organized in consortia comprising multiple interacting populations that cooperate to achieve a common goal [1]. This self-organization increases the survival fitness of the community and, by specialization and division of labor, reduces the burden on each population while increasing the overall efficiency of the consortium [2]. Over the past decade the translation of these advantages to synthetically engineered biological systems has attracted increasing interest [3] as it is an effective solution for the efficient production of complex chemical compounds [4]– [6].

In addition to the design of the phenotype of each population, the realization of functional microbial consortia requires stable, long-term coexistence between the different populations therein. This is a challenging goal to achieve as each population, engineered with different genetic regulatory networks, is burdened by a different metabolic load, resulting in heterogeneous growth rates. Left uncontrolled, this imbalance will ultimately lead to the extinction of one or more populations, disrupting the proper functioning of the consortium. Thus, the key issue of regulating the microbial community’s composition, known as composition, or ratiometric, control, is essential for creating consortia capable of reliably performing some desired function [7].

A promising scalable approach to co-culturing multiple bacterial populations involves using bioreactors to control both the density and composition of the consortium. Various strategies for managing coexistence within a single bioreactor have been explored. In [8] it was employed a deep reinforcement learning algorithm to adjust the growth media composition, ensuring coexistence of two microbial populations dependent on different substrates. Similarly, studies in [9], [10] showed that modifying the dilution rate can support robust coexistence of two *complementary strains*. These approaches presume that growth conditions can be selectively altered to favor one population over another. However, it is not always feasible to create such conditions, as the growth properties, determined by the cellular resources consumed up to a given time, may remain fixed. In cocultures, this results in the population that consumes more resources growing slower and potentially facing extinction. In a single bioreactor, this is impossible or cumbersome to avoid even when using a sophisticated feedback control action, because of the competitive exclusion principle [11], which from a dynamical systems standpoint translates into the system being uncontrollable. Therefore, a more general methodology is needed to guarantee long-term coexistence of different microbial species in a bioreactor.

In this Letter, we propose an experimental platform endowed with a control strategy for robust regulation of both the density and the composition of a microbial consortium made of two non-complementary strains (one population persistently growing faster than the other). Specifically, we propose to use two communicating bioreactors (turbidostats) (see Fig. 1); one hosting a mixture of both populations, while the other being used as a reservoir where the slower population is grown alone. By using as control inputs the dilution rate *D*_1_ (inlet volumetric flow/volume) associated with fresh media in the first reactor in combination with the dilution rate *D*_2_, at which the slower population from the reservoir (kept at a reference density therein by means of the dilution rate *D*_0_) is added to the mixing chamber, our strategy can guarantee coexistence of the two populations at a desired ratio.

**Fig. 1.**
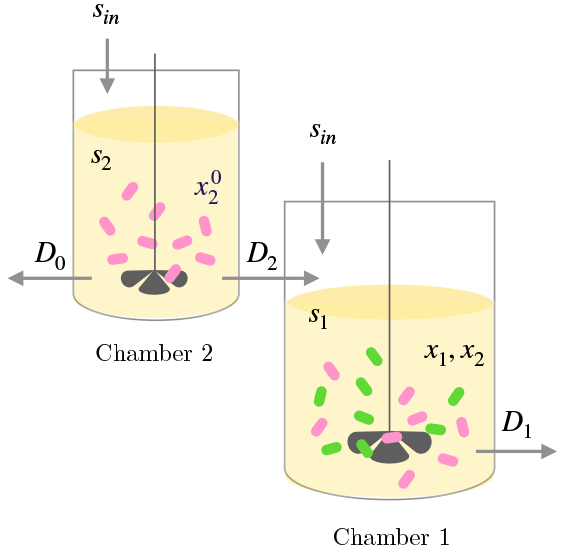
Schematic representation of the two-reactor architecture. The slower species (species 2, in pink) is grown separately in chamber 2 and added at rate *D*_2_ in chamber 1, where it is mixed with species 1 (in green). Species 2 is regulated at the desired concentration 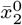 in chamber 2 by means of the dilution rate *D*_0_. The dilution rate *D*_1_ is used in combination with *D*_2_ to keep the two species at the desired concentrations in chamber 1.

In what follows, we introduce in Section II a mathematical model that describes the growth dynamics of cells within each reactor and analyze its asymptotic behavior. Section III outlines the statement of the control problem, which is addressed in Section IV using an open-loop strategy. A closed-loop control law is detailed in Section V. Both strategies are validated *in silico* in Section VI using a model parametrized with real experimental data from two Chi.Bio turbidostats [12]. The proposed approach is versatile, suitable for a broad range of non-complementary strains growing on the same substrate, and does not require genetic engineering interventions.

## II. Mathematical Model

We consider two microbial species growing as a continuous culture in a chemostat (labeled as chamber 1 in Fig. 1), with a reservoir (chamber 2 in Fig. 1) in which one species grows separately. The two species are assumed to be independent, that is, they grow on the same substrate but they do not directly condition the growth rate of each other. The mathematical model of the former chemostat with two species and two inputs can be obtained by applying straightforward balance of mass law as in [13], yielding

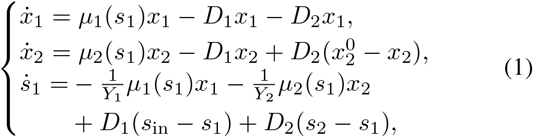

where the variables *x*_1_, *x*_2_ ∈ ℝ_≥0_ and *s*_1_, *s*_2_ ∈ [0, *s*_in_] denote the concentrations of the biomass of species 1 and 2 and of the substrate in chamber 1 and 2, respectively. The mathematical model of the reservoir (chamber 2) can be written as

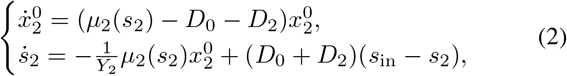

where 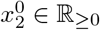 denotes the concentration of the biomass of species 2 in chamber 2. Moreover, *µ*_*i*_(·) is the growth rates of species *i* (defined below), *Y*_*i*_ is the yield coefficient, assumed without loss of generality to be unitary for both species, and *s*_in_ is the concentration of the substrate in the inlet flows. The control inputs *D*_*i*_ are the dilution rates defined as the ratio between the inlet flow rate and the culture volume. Specifically, the control inputs *D*_1_(*t*) : ℝ_≥0_ ↦ [*D*_1,min_, *D*_1,max_], with *D*_1,min_ ≥ 0, and *D*_0_(*t*) : ℝ_≥0_ ↦ [*D*_0,min_, *D*_1,max_], with *D*_0,min_ > 0, dilute the concentrations in chambers 1 and 2, respectively, by injecting fresh substrate at concentration *s*_in_, while the input *D*_2_(*t*) : ℝ_≥0_ ↦ [0, *D*_2,max_] is the dilution rate due to the inlet flow of media from chamber 2 to chamber 1. The dilution rates are assumed to be the same for both species, that is, the culture is well-mixed and mortality and attachment of the bacteria are neglected [10]. The dynamics of systems (1)-(2) is characterized by the growth rate functions *µ*_*i*_(·) : [0, *s*_in_] ↦ ℝ_≥0_, that are typically assumed to be differentiable, strictly increasing, and such that *µ*_*i*_(0) = 0, for *i* = 1, 2, corresponding to the substrate not having any inhibitory effect at high concentrations [14]. The most common analytical growth rate model, that we also consider here, is the so-called Monod law [14]

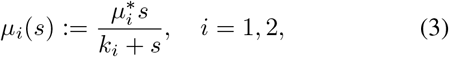

where 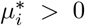 is the maximum growth rate of species *i* and *k*_*i*_ > 0 is a Michaelis-Menten constant. When the two growth functions intersect more than once, it is possible to design some control input *D*_1_(*t*) (setting *D*_2_ = 0) such that both species survive [9], [10], [15]. On the contrary, when it holds that

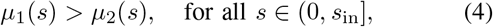

the growth functions intersect only at 0 and the two species are said to be non-complementary [10], i.e. one species always outgrowing the other. Therefore, in this condition, from the Competitive Exclusion Principle (CEP) [16], it follows that when inflow from the additional reservoir is not available, i.e., *D*_2_ = 0, no control input *D*_1_(*t*) exists such that more than one strain survives at steady state. However, in our architecture (Fig. 1) coexistence is still made possible by a controlled injection into the chemostat (chamber 1) of some *extra* biomass of the slower species taken from the reservoir (chamber 2) by means of the additional control input *D*_2_(*t*).

### A. Reduced system

Under the assumption that the sum of the control inputs *D*_0_(*t*) and *D*_2_(*t*) is persistently exciting, that is, such that (*D*_0_(*t*) + *D*_2_(*t*)) > 0, ∀*t* ≥ 0, the solutions of system (2) are attracted to the invariant subspace 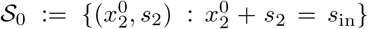. This can be demonstrated using invariant set theory, as illustrated in [17], similarly to the approach used in [15, Proposition 1] (see also Appendix for further details). Thus, if the initial condition 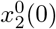 belongs to 𝒮_0_, the dynamics of (2) on 𝒮_0_ can be reduced to

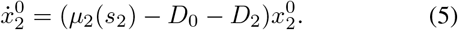

Likewise, under the assumption that the sum of *D*_1_(*t*) and *D*_2_(*t*) is persistently exciting, it can be proved using similar arguments that the solutions of system (1) are attracted to the invariant subspace 𝒮 := {(*x*_1_, *x*_2_, *s*_1_) : *x*_1_ + *x*_2_ + *s*_1_ = *s*_in_}. Thus, solutions rooted therein can be obtained from the reduced order model

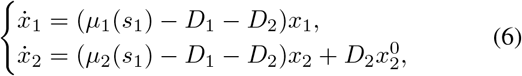

defined on the invariant set 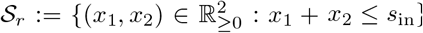.

### B. Stability analysis of the reservoir model

Depending on the value of (*D*_0_ + *D*_2_), system (5) can have one or two equilibria. One equilibrium is always at 0, while the other is 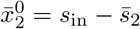, with

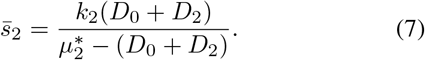

The point 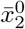 is admissible and stable for 0 < (*D*_0_ + *D*_2_) < *µ*_2_(*s*_in_). It becomes non-admissible, exchanging its stability with the point at the origin, when (*D*_0_ + *D*_2_) = *µ*_2_(*s*_in_), through a transcritical bifurcation.

### C. Stability analysis of the chemostat model

System (6) has a richer dynamics depending on the values of *D*_1_ and *D*_2_.

1. *D*_1_ > 0, *D*_2_ > 0: When 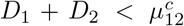, with 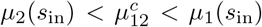, there exists a stable equilibrium point in *S*_*r*_ such that both species coexist. This point is located at the intersection between the *x*_1_-nullcline 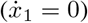 and the *x*_2_-nullcline 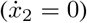, obtained by solving the system of equations

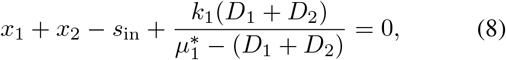

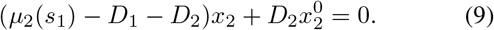

Moreover, there exists another equilibrium point, that is unstable, lying on the *x*_2_-axis (*x*_1_ = 0) at the intersection with (9). When *D*_1_ + *D*_2_ is above some critical value 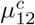, the coexistence point collides with the unstable point on the *x*_2_-axis, exchanging stability via a transcritical bifurcation, and ceases to be admissible. At this point only species 2 survives, because the total dilution rate in the bioreactor is too high for species 1 to survive. However, note that the complete extinction of species 1 is not possible in practice, as experimentally some cells will not be flushed out. Due to the nonlinearity of (9), the analytical expression of 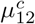 is cumbersome and is omitted here for the sake of brevity.
2. *D*_1_ ≈ 0, *D*_2_ > 0: The dynamics is similar to the previous case, that is, there exists a stable equilibrium point associated with coexistence whose position depends only on *D*_2_. However, when *D*_2_ > *µ*_1_(*s*_in_), that is, the dilution rate from the reservoir is higher than the maximum growth rate of species 1, the faster species is flushed out from the bioreactor 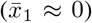 and as *D*_2_ increases the concentration of species 2 reaches asymptotically the same value as in the reservoir (i.e.,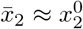).
3. *D*_1_ > 0, *D*_2_ = 0: Model (6) becomes the same model described in [9], and as *D*_1_ varies from 0 to *D*_1,max_ > *µ*_1_(*s*_in_) we retrieve the same dynamics depicted in the phase portraits shown in Fig. 2 from [9] (namely cases I, IV, V, VI).
4. *D*_1_ = 0, *D*_2_ = 0: All solutions converge to the attractive equilibrium set ε_0_ := {(*x*_1_, *x*_2_) ∈ 𝒮_*r*_ : *x*_2_ = −*x*_1_ + *s*_in_} corresponding to the biomass being in starvation and therefore it is not an admissible working condition in continuous culture (see case I in Fig. 2 from [9]).

**Fig. 2.**
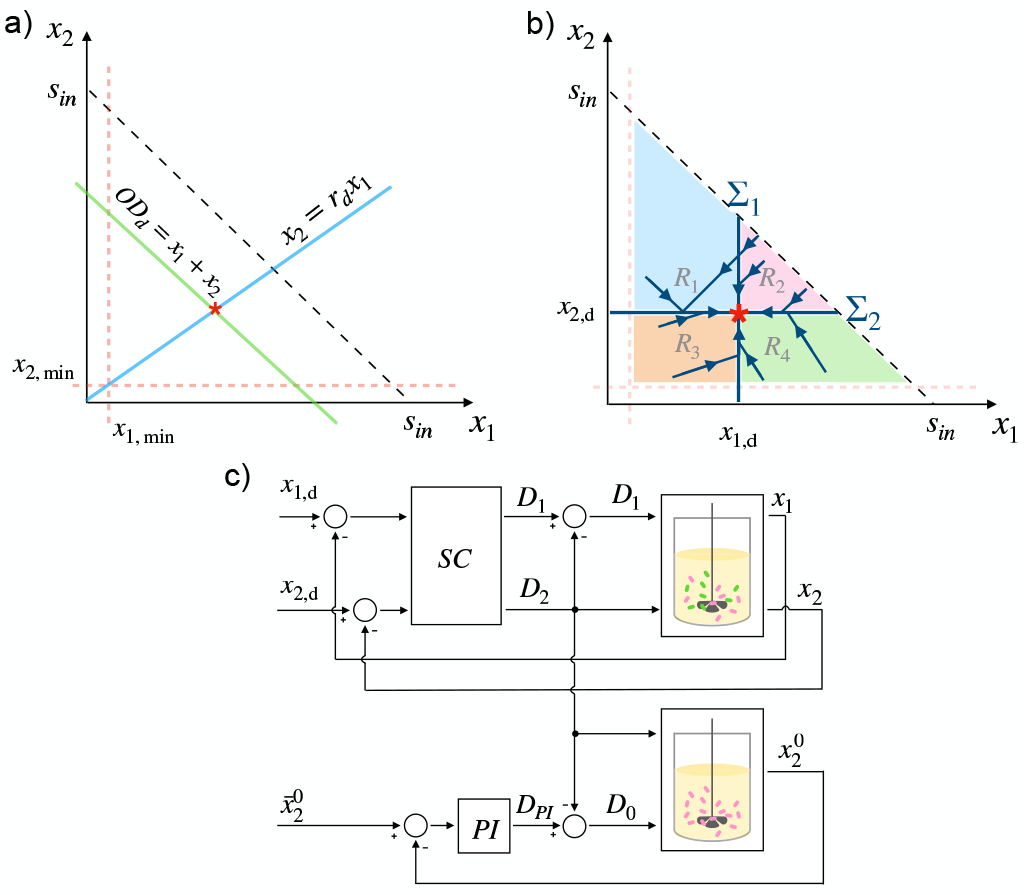
a) Graphical representation of the control objective. The green and blue lines represent the sets where *x*_1_ + *x*_2_ = *OD*_d_ and *x*_2_ = *r*_d_*x*_1_, respectively. The black dashed line delimits the invariant set 𝒮_*r*_. Finally, the orange dashed lines indicate the safety bounds for each species (i.e. *x*_1,min_, *x*_2,min_). b) Graphical representation of the closed-loop control strategy. 𝒮_*r*_ is divided in four regions (*R*_1_, *R*_2_, *R*_3_, *R*_4_) by the switching surfaces Σ_1_ and Σ_2_ (blue lines). The control actions *D*_1_ and *D*_2_ steer the trajectory of the system (blue arrows) onto *x*_d_ (red star). c) Block diagram of the closed-loop control architecture. The switching controller *SC* regulates the concentrations of the biomasses *x*_1_ and *x*_2_ in chamber 1 to the desired levels *x*_1,d_ and *x*_2,d_, respectively. The control input *D*_0_, regulating 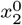 in chamber 2 to the desired level 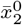, is computed by a PI controller with a feed-forward disturbance compensation.

## III. Control Problem

Given the two-chambers bioreactor architecture, depicted in Fig. 1, with two *non-complementary* microbial species, i.e. such that condition (4) holds, whose mathematical model and dynamics have been described in Sec. II, our control objective is to control the dilution rates such that at steady state the two species coexist and their concentrations are regulated at a desired ratio. That is, we want to solve the following problem.

### Control problem

Design some control inputs *D*_0_(*t*), *D*_1_(*t*), and *D*_2_(*t*) for system (5)-(6), under condition (4), such that:

1. *coexistence* of the two species is guaranteed for all time, that is, *x*_*i*_(*t*) ≥ *x*_*i*,min_, ∀*t* > 0, *i* = 1, 2, where *x*_*i*,min_ > 0 is some safety value to avoid extinction of species *i*;
2. the *total biomass* is regulated to a desired value to guarantee efficient utilization of the resources, that is,

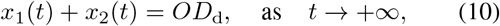

where *OD*_d_ is the desired optical density in chamber 1, assumed for the sake of simplicity to be an exact proxy of the total biomass therein;
3. the *ratio x*_2_*/x*_1_ between the concentration of the two species in chamber 1 is robustly regulated at steady state to a desired value *r*_d_, that is,

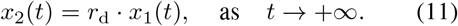

Notice that the above control problem corresponds to requiring that all solutions of system (5)-(6), starting from any initial conditions in 𝒮_0_ ∪ 𝒮_*r*_, converge to the point of intersection between the two curves defined in (10) and (11) (Fig. 2.a), that is, to the point

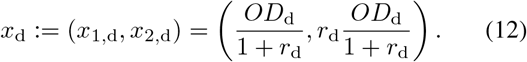

In the following, we assume that the concentrations *x*_1_, *x*_2_ and 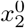 are either directly or indirectly measurable, for example via fluorescent reporters or state observers.

## IV. Open-Loop Control

A simple open-loop controller that solves the control problem described above can be designed by exploiting the dynamics of the system described in Sec. II, that is, by choosing constant values 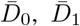 and 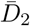 so as to make the nullclines (8) and (9) intersect at the desired point (12). Specifically, setting 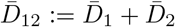 by using (8) and (10) we obtain

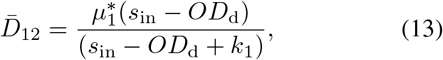

Moreover, by using (9) and (12) we get

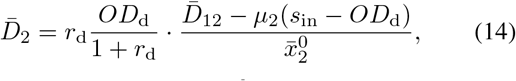

where the steady-state value 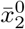 in the reservoir can be reached by setting the control input *D*_0_ to 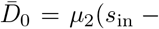 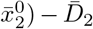, obtained from setting the first term in (5) to zero. Obviously, we also have that 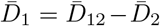. Further details about the derivation of (13)-(14) are reported in Appendix.

Although this feed-forward control action is simple and solves the control problem in nominal conditions, as we will see later in Sec. VI, it neither guarantees robustness to perturbation nor faster convergence. The coexistence requirement, as defined in Sec. III, is fulfilled by the open-loop controller provided that the initial condition (*x*_1_(0), *x*_2_(0)) is chosen beneath the intersection between the nullcline (8) and the constraint *x*_1_ ≥ *x*_1,min_.

## V. Closed-Loop Control

In biological applications robustness is as important as precision in the regulation and open-loop strategies cannot often reject external perturbations and uncertainties in the model parameters. Therefore, we propose next a closed-loop switching controller that can globally and robustly guarantee regulation of the desired density and composition.

The controller is designed by defining two switching surfaces, Σ_1_ := {(*x*_1_, *x*_2_) ∈ 𝒮_*r*_ : *x*_1_ = *x*_1,d_} and Σ_2_ := {(*x*_1_, *x*_2_) ∈ *S*_*r*_ : *x*_2_ = *x*_2,d_}, which divide the domain 𝒮_*r*_ into four regions (see Fig. 2.b), *R*_1_ where *x*_1_ < *x*_1,d_ and *x*_2_ > *x*_2,d_, *R*_2_ where *x*_1_ > *x*_1,d_ and *x*_2_ > *x*_2,d_, *R*_3_ where *x*_1_ < *x*_1,d_ and *x*_2_ < *x*_2,d_, and *R*_4_ where *x*_1_ > *x*_1,d_ and *x*_2_ < *x*_2,d_. We denote by Σ_*ij*_ the boundary between regions *R*_*i*_ and *R*_*j*_.

### Theorem 1

Choosing 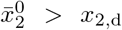 as set-point for the reservoir, and the switching control inputs as

- 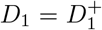 and *D*_2_ = 0, with 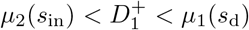, with *s*_d_ := *s*_in_ − *OD*_d_, when *x* ∈ *R*_1_,
- 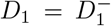 and *D*_2_ = 0, with 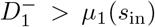, when *x* ∈ *R*_2_, so that the bioreactor is in flush-out mode,
- *D*_1_ = *D*_2_ = 0 when *x* ∈ *R*_3_, so that way the biomass is left to grow freely,
- *D*_1_ = 0 and 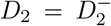, with 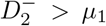, when *x* ∈ *R*_4_,

the setpoint *x*_d_, as defined in (12), is attractive for all solutions of system (6) in 𝒮_*r*_.

*Proof:* This can be proved by showing that the intersection of the two switching surfaces, Σ_1_ ∩ Σ_2_ is attracting. Let *n*_1_ = [1 0]^*T*^, *n*_2_ = [0 1]^*T*^ be the normal vectors to Σ_1_ and Σ_2_, respectively, and *f*_1_(*x*), *f*_2_(*x*), *f*_3_(*x*), and *f*_4_(*x*) the vector fields of system (6) in each of the four regions determined by our choice of the switching control inputs, that is,

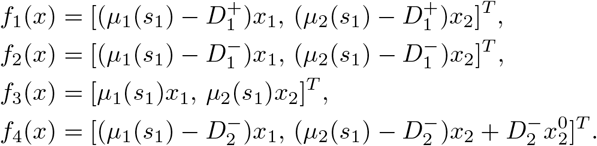

The boundary Σ_13_ is a stable sliding region because 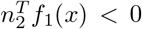 and 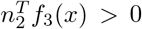, since 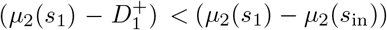 and *µ*_2_(*s*_1_) > 0, respectively. Moreover, solutions sliding on Σ_13_ are attracted towards *x*_d_ because therein 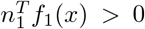 and 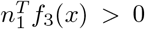, since, if 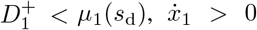 when *x*_1_ < *x*_1,d_. This is because the intersection between the nullcline (8) and Σ_2_ lies above *x*_d_, and *µ*_*s*_(*s*_1_) > 0, respectively. Similarly, Σ_34_ is a stable sliding region because 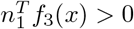 and 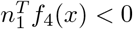, since 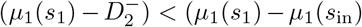. Moreover, solutions on Σ_34_ are attracted towards *x*_d_ because therein 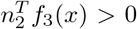 and 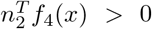, since 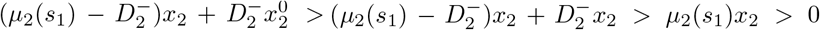, where we used 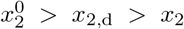. Likewise, Σ_24_ is a stable sliding region because 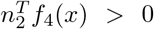 and 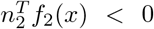, since 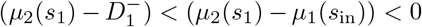. Solutions on Σ_24_ are attracted towards *x*_d_ because therein 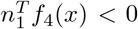 and 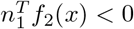, since 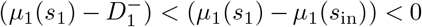. Finally, the boundary Σ_12_ has always a crossing region, in which solutions from *R*_2_ cross the boundary because 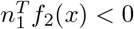 while 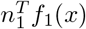 changes sign at the intersection of Σ_12_ with the nullcline (8). However, all crossing solutions will then converge onto Σ_13_ since 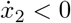, and therefore they will move towards *x*_d_, as proved above. This concludes the proof. ◼

Although as stated earlier, complete extinction of species 1 is not possible, some trajectories starting within *R*_1_ ⊂ 𝒮_*r*_ may still violate the coexistence bounds defined in Sec. III. The set of initial conditions (*x*_1_(0), *x*_2_(0)) ∈ *R*_1_ yielding trajectories that temporarily violate those bounds can be removed by choosing 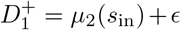, with ϵ > 0 arbitrarily small, so that the nullcline (8) intersects at the uppermost point the line *x*_1_ = *x*_1,min_. Therefore, initial conditions must be chosen so that 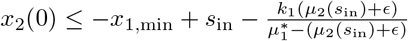.

The controller implemented in the reservoir (chamber 2) to regulate 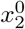 is a PI controller with a disturbance compensation. Specifically, the dilution rate *D*_0_ in (5) is chosen as *D*_0_ = *D*_*PI*_ − *D*_2_, where *D*_*PI*_ is an action computed using the same proportional-integral controller presented in [12] for the regulation of the turbidity. This controller ensures that the error 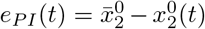 converges to zero, so that the density of species 2 in the reservoir 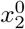 is regulated to the desired value 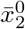.

## VI. In silico validation

We validated the designed controllers *in silico* to assess their performance, stability, and robustness. Specifically, we simulated the experimental platform shown in Fig. 1 implemented using two Chi.Bio reactors [12] and including the technological constraints imposed by the experimental platform. Each reactor measures the density of the biomasses at every minute by measuring the turbidity of the liquid (i.e. the optical density) and updates the duty-cycle of the peristaltic pumps that dilute the cultures with fresh media. The dilution rates are saturated to 0.065 min^−1^ for safety reasons. In all control experiments we set the desired reference signals to *OD*_d_ = 0.7 and *r*_d_ = 1. The parameters of the simulated set-up were parametrized from real data as explained below.

### A. Model parametrization

For the identification of the parameters in system (6), we conducted open-loop characterization experiments using the commercially available reactor Chi.Bio [12]. Specifically, we characterized the growth dynamics of two *E. coli* strains embedding two different implementations of a genetic toggle-switch, presented in [18] and [19]. In each experiment, using a single Chi.Bio reactor, we grew a single strain in LB media supplemented with 50 *µg/µL* kanamycin and 1 *mM* isopropyl *β*-D-1-thiogalactopyranoside (IPTG), at 37°*C*, shaking at half the maximum speed available on the platform. During these experiments, the culture was diluted using the maximum available dilution rate until the optical density dropped below 0.3. Then, cells underwent a recovery phase where the dilution rate was kept close to zero for 60 minutes. Finally, the cells’ growth was excited changing the dilution rates every 15 to 30 minutes for 1 hour (Fig. 3).

**Fig. 3.**
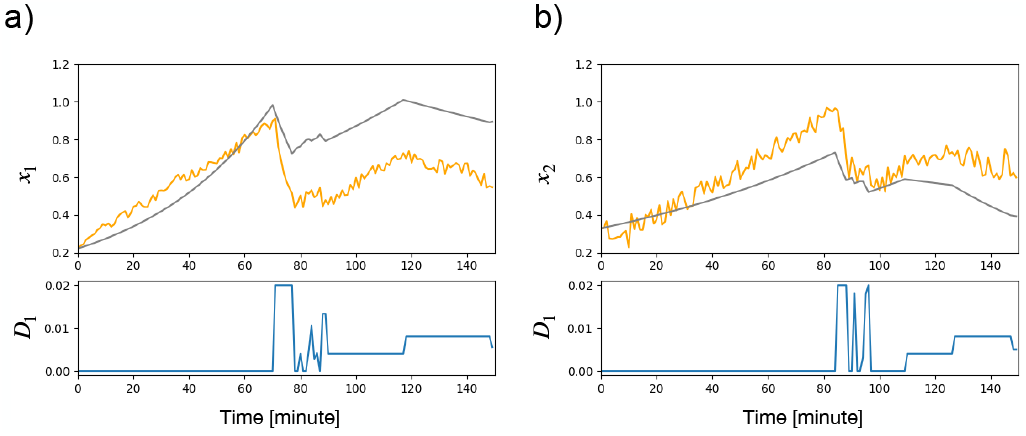
Comparison between model predictions and the experimental data obtained from open-loop characterization experiments conducted using the Chi.Bio reactor. Two different *E. coli* strains implementing genetic toggle-switches from [18] (panel a) and [19] (panel b) were used in the experiments. The top panels depict the temporal evolution of the biomass in the *in vivo* experiment (orange) and in the numerical simulation (gray). The bottom panels show the inputs applied to the bioreactors. The experimental data are collected using a single reactor (i.e. setting *D*_0_ = *D*_2_ = 0).

Using Genetic Algorithm (GA) optimization (as implemented in [20]), we estimated the growth parameters for each strain to be: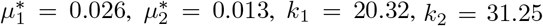, and *s*_in_ = 88.75. The ranges for the 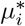 were set according to typical values in literature [19], while the other parameters’ ranges were obtained heuristically. The intervals were set as follows: [0.001, 0.05] for 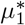 and 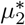, [0.01, 100] for *k*_1_ and *k*_2_, [1, 100] for *s*_in_ to start the GA optimization. The total number of iterations was 200, and the initial population of the GA algorithm was set to be 150 individuals. In Fig. 3 the model’s predictions are depicted alongside the experimental data, demonstrating the model ability to capture quantitatively the growth dynamics of the microbial species under varying conditions. The data used for validation were different from those employed for the parameters identification. The quality of the predictions was quantified using the Root Mean Squared Error (RMSE), which proved to be low for the estimation of both populations’ trajectories (i.e. RMSE = 3.18 for species 1 and RMSE = 1.73 for species 2) and comparable with the RMSE of the identification data set (i.e. RMSE = 1.86 for species 1 and RMSE = 1.09 for species 2).

### B. Control validation

Firstly, we simulated model (1)-(2) under the action of the open-loop controller, described in Sec. IV (with dilution rates equal to 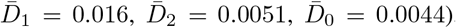), including also the technological constraints described above, observing that the desired density and configuration were reached asymptotically (see Fig. 4.a). However, the convergence time was excessively long, namely taking 97 · 10^3^ min and 116 · 10^3^ min to reach the desired optical density and ratio, respectively, rendering this strategy unfeasible in practice for *in vivo* implementation.

**Fig. 4.**
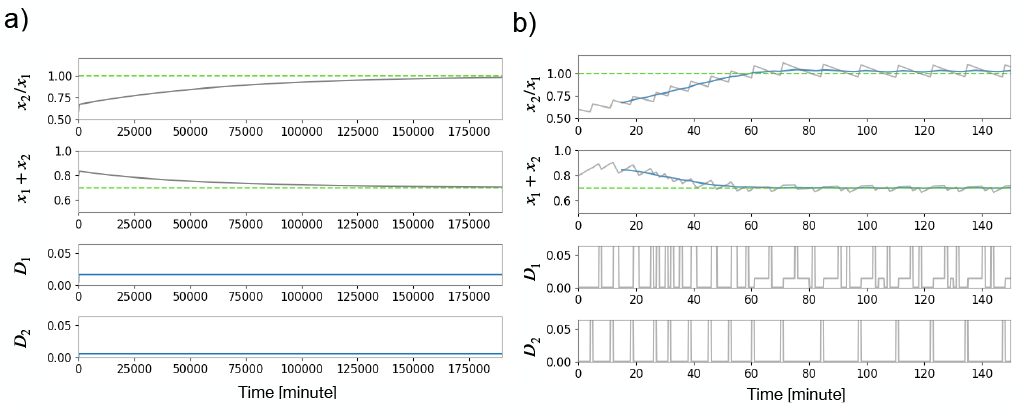
*In silico* validation of the open-loop (a) and closed-loop (b) control strategies. The top panels show the evolution in time of *x*_2_*/x*_1_ and *x*_1_ +*x*_2_, depicted as gray solid lines, with their reference values (dashed green lines) and, for those in (b), the average trajectory over a time window of 30 minutes (blue solid line). The reference values are set as *OD*_d_ = 0.7 and *r*_d_ = 1. The bottom panels show the inputs applied to the first chamber. The control parameters of the closed-loop controller are set to 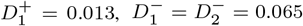.

Instead, as shown in Fig. 4.b the switching control strategy developed in Sec. V was able to guarantee a much faster settling time (at 5%), namely, 40.5 min and 52.6 min for the optical density and the ratio between the two species, respectively, at the cost of a relatively small residual error at steady state. We quantified the relative residual errors by evaluating the average deviation from the desired values in the last 60 minutes of the experiments, that is, 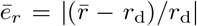, where 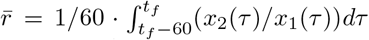, where *t*_*f*_ = 150 min, and similarly for *ē*_*OD*_, obtaining 0.026 and 0.0004, respectively.

Finally, we assessed the robustness of the proposed switched controller against variations in the parameters of the growth functions *µ*_*i*_(*s*), *i* = 1, 2. Specifically, the values of the parameters 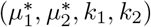 were randomly drawn from normal distributions, each centered on their nominal value, say *η*, with a standard deviation equal to *σ* = *CV* · *η*, where *CV* is the coefficient of variation. For each value of *CV*, taken in the interval [0.05, 0.3], we evaluated the mean and standard deviation of the residual errors 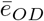 and 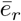, over 100 simulations, each computed with different values of the parameters 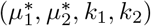. The results of this analysis, reported in Fig. 5, highlight that, even when a consistent mismatch is present between the identified parameters and their true value (*CV* = 30%), the average steady-state residual errors remain below 0.05 for the ratio and 0.01 for the optical density.

**Fig. 5.**
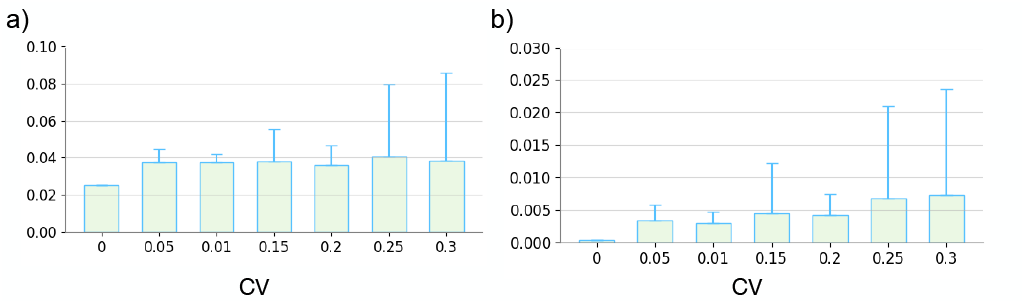
Robustness analysis of the closed-loop controller. Mean (bar height) and standard deviation (whisker) of *ē*_*r*_ (panel a) and *ē*_*OD*_ (panel b), for increasing value of the coefficient of variation *CV*. Each bar is obtained by averaging 100 simulations each computed with different values of parameters 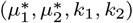.

## VII. CONCLUSIONS

We introduced a dual-reactor architecture to steer the overall density and the composition of a two-strain consortium. We analyzed its open-loop dynamics and then formulated two tailored control strategies. These strategies were validated *in silico* with model parameters identified experimentally. Our approach enables long-term coexistence within a consortium of non-complementary cell populations without the integration of additional synthetic circuits into the cells. This capability is crucial for supporting the simultaneous growth of different populations in a consortium, potentially across varied species such as bacteria and yeasts. We are currently implementing a prototype of the proposed architecture, shown in Fig. 1, featuring two Chi.Bio reactors. Ongoing *in vivo* experiments aim to further validate the effectiveness of our control strategies, with results to be detailed in forthcoming publications.

## Acknowledgements

The authors acknowledge support from the Telethon Institute of Genetics and Medicine (TIGEM) in Naples, Italy for their invaluable support and collaboration throughout this research project. DF and GR wish to acknowledge financial support by the European Union - Next Generation EU, under PRIN 2022 PNRR, Project “Control of smart microbial communities for wastewater treatment”. MdB wishes to acknowledge financial support by the Italian Ministry of Health under PNRR PNC, Project “Digital Health Solutions in Community Medicine (DHEAL-COM)” – Project Code PNC-E3-2022-23683267 PNC – HLS – DH.

## APPENDIX

### A. Attractivity and invariance of 𝒮_0_ and 𝒮

Convergence of the solutions of system (1) and (2) to the subspaces 𝒮_0_ and 𝒮, respectively, can be proved my means of the LaSalle’s Invariance Principle, similarly as done in [15]. Specifically, concerning the subspace 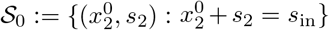, consider the function 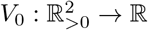 defined as

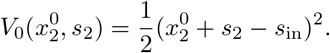

Taking the time derivative and using the dynamics in (2), we get 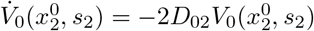, where *D*_02_ = *D*_0_ + *D*_2_. Now, provided that *D*_0_(*t*) + *D*_2_(*t*) is persistently exciting, we have that 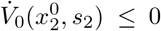 for all 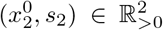. By LaSalle’s Invariance Principle, all solutions converge to the largest invariant set *M* contained in the set 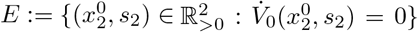. Since 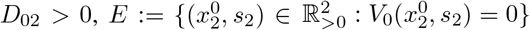. Therefore, all solutions 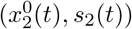 starting in 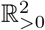 asymptotically converge to 𝒮_0_.

Likewise, concerning the subspace 𝒮 := {(*x*_1_, *x*_2_, *s*_1_) : *x*_1_ +*x*_2_ +*s*_1_ = *s*_in_}, we can employ the function 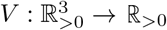 defined as

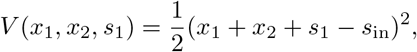

whose time derivative is 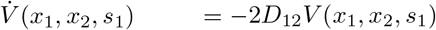, where we assumed that 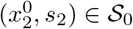, and *D*_12_ = *D*_1_ + *D*_2_. Following similar steps as above, we can conclude that all solutions (*x*_1_(*t*), *x*_2_(*t*), *s*_1_(*t*)) starting in 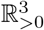 converge to 𝒮.

### B. Open-loop control actions

Regarding the derivation of the open-loop control actions (13) and (14), the former is obtained by substituting (10) into (8), getting

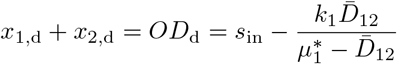

Where 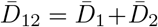. Then, by solving for 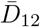 one obtains (13).

For what concerns (14), substituting (10) into (9) we get

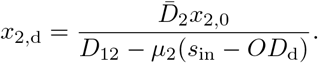

Now, using the expression of *µ*_2_(·) evaluated at the point *s*_1_ = *s*_in_ − *OD*_d_, that is,

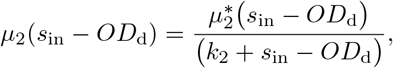

and that of *x*_2,d_, obtained by combining (10) and (11),

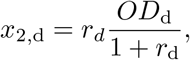

we get

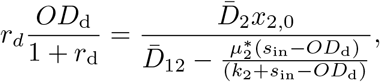

that, by solving for 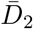, finally gives (14).

## Notes

### Competing Interest Statement

The authors have declared no competing interest.

### Summary of Updates

Major: we have revised the closed-loop control strategy and we added details concerning the derivation of the Open-Loop Control inputs and the attractivity and invariance of the subsets of the reduced system (See. Appendix). Minor: we have changed the structure of the section to improve readability.

